## Supplemental Figure S1 for "Reporting and Misreporting of Sex Differences in the Biological Sciences"

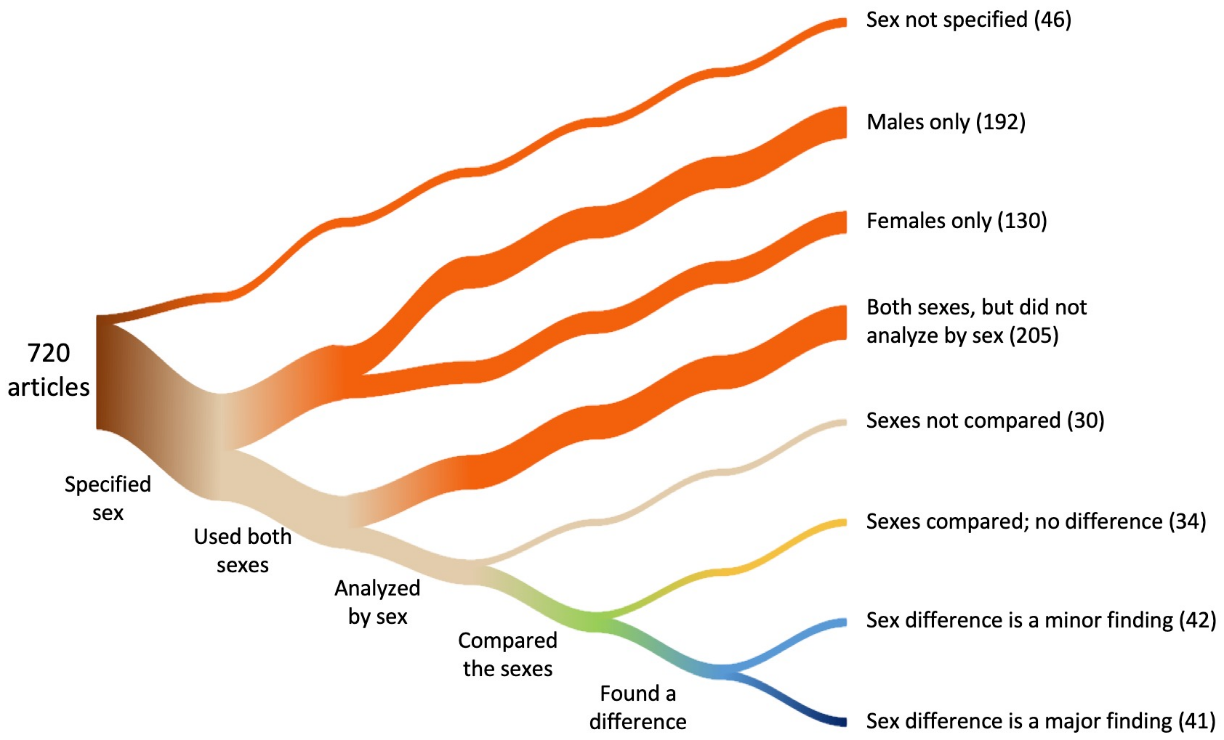


**Fig. S1. River plot showing our findings in the larger context of the study by Woitowich et al. (2020).** The width of each stream is proportional to the number of articles represented in that stream. Starting with 720 articles, Woitowich et al. asked whether sex was specified, whether only one or both sexes were included, and whether the data were analyzed by sex. In this study, we began with the 147 articles in which data were analyzed by sex and asked further whether the sexes were compared and what was found (lower four branches; see Main Text Fig. 1A).
